# BMDD: A Probabilistic Framework for Accurate Imputation of Zero-inflated Microbiome Sequencing Data

**DOI:** 10.1101/2025.05.08.652808

**Authors:** Huijuan Zhou, Jun Chen, Xianyang Zhang

**Author notes:** Address correspondence to Jun Chen and Xianyang Zhang.

## Abstract

Microbiome sequencing data are inherently sparse and compositional, with excessive zeros arising from biological absence or insufficient sampling. These zeros pose significant challenges for downstream analyses, particularly those that require log-transformation. We introduce BMDD (BiModal Dirichlet Distribution), a novel probabilistic modeling framework for accurate imputation of microbiome sequencing data. Unlike existing imputation approaches that assume unimodal abundance, BMDD captures the bimodal abundance distribution of the taxa via a mixture of Dirichlet priors. It uses variational inference and a scalable expectation-maximization algorithm for efficient imputation. Through simulations and real microbiome datasets, we demonstrate that BMDD outperforms competing methods in reconstructing true abundances and improves the performance of differential abundance analysis. Through multiple posterior samples, BMDD enables robust inference by accounting for uncertainty in zero imputation. Our method offers a principled and computationally efficient solution for analyzing high-dimensional, zero-inflated microbiome sequencing data and is broadly applicable in microbial biomarker discovery and host-microbiome interaction studies. BMDD is available at: https://github.com/zhouhj1994/BMDD.

**Author Summary:** Understanding the microbes living in and on our bodies—the microbiome—relies on analyzing complex sequencing data. However, these data often contain many zeros, either because a microbe is truly absent or simply missed due to insufficient sampling. These missing values make it hard to accurately analyze microbial patterns and identify important differences between groups, especially for methods that work on a log scale. To address this, we developed a new method called BMDD that uses a more realistic model to impute the zeros. Unlike existing tools that assume each microbe follows an unimodal abundance distribution, BMDD allows for microbes to follow a bimodal distribution, so they could behave differently in different conditions. It provides not just a single guess, but a range of possible values to better reflect the uncertainty. Our testing shows that BMDD more accurately recovers the true microbial profiles and improves the ability to detect meaningful differences between groups. This method can help researchers gain clearer insights into how the microbiome affects health and disease.

## 1 Introduction

The human microbiome has been emerging as an important player in health and disease (Kashyap et al. 2017, Gevers et al. 2014, Jostins et al. 2012, Kostic et al. 2012, Qin et al. 2012, Scher et al. 2013, Qin et al. 2014). For example, immune maturation and modulation (Ahern et al. 2014, Geva-Zatorsky et al. 2017), inflammatory cytokine production (Schirmer et al. 2016), host serum metabolome and insulin level (Pedersen et al. 2016), and host gene regulation (Fellows et al. 2018) have all been shown to be linked to the human microbiome. Complementary to existing omics data, microbiome data offers an additional perspective on human health (Kashyap et al. 2017). The dysbiosis of the human microbiome has been implicated in numerous human diseases (Young 2017).

Next-generation sequencing allows for the determination of the microbiome composition by direct microbial DNA sequencing (Kuczynski et al. 2012). Microbiome taxonomic abundance data, usually in the form of a count table of detected taxa, are typically over-dispersed and sparse with many zeros. Zeros could be due to either the physical absence of the taxa or their low abundance so that the sequencing machine cannot detect them reliably. Excessive zeros pose many statistical challenges for microbiome data analysis (Yang & Chen 2022) and are particularly problematic for logarithmic scale analysis. The traditional approach involves adding a pseudocount, such as 1 or 0.5, to all counts before normalizing the data into relative abundances. Downstream statistical analyses are then based on the relative abundance data (a.k.a. compositional data), which sum up to one for each sample. Adding a pseudocount has been shown to have detrimental consequences in certain contexts (Brill et al. 2022).

The pseudocount approach, as a naive method to impute the zeros, does not fully exploit the information in the data and thus is far from optimal. An efficient zero-imputation approach should be able to tap into the correlation structure and the distributional characteristics of the data. To improve over the pseudocount approach, various zero-imputation techniques have been proposed for zero-inflated count data, including single-cell RNA sequencing (scRNA-seq) and microbiome sequencing data. One popular approach is to assume an underlying true abundance or expression matrix and estimate it using Bayesian methods. Examples of such methods include SAVER (Huang et al. 2018) for scRNA-seq data and mbDenoise (Zeng et al. 2022) for microbiome data. SAVER models the count using a Poisson distribution, where the mean is the product of the true gene expression level and a normalization factor. A gamma prior is imposed on this true expression level. The hyperparameters in the priors are empirically estimated by fitting linear regression models between each gene and other genes in the same cell. On the other hand, mbDenoise uses zero-inflated negative binomial (ZINB) distributions to model taxon abundances. The mean of the negative binomial distribution is represented by the low-rank approximation of the count matrix. The posterior mean of the ZINB model is estimated using variational approximation and is used to recover the true abundance level. Another approach to microbiome data imputation, mbImpute (Jiang et al. 2021), assumes that each taxon’s abundance follows a mixture model: a gamma distribution for missing abundances requiring imputation and a log-normal distribution for actual (non-missing) abundances. For abundance values identified as non-missing, mbImpute fits linear models that combine information from similar samples, similar taxa, and sample covariates, then imputes the missing values by the fitted values. The authors of mbImpute also proposed a method called scImpute for imputing scRNA-seq data employing principles similar to those used in mbImpute (Li & Li 2018). ALRA is another method for imputing scRNA-seq data (Linderman et al. 2022), which uses singular value decomposition for low-rank approximation of the expression matrix. The rank is determined by identifying noise singular values, and data below a specified quantile after low-rank approximation is thresholded to preserve biological zeros.

Existing imputation methods either explicitly or implicitly assume a feature-wise unimodal distribution of the nonzero proportions/counts, which may be too restrictive for real microbiome abundance data. Here, we propose a more general Bayesian approach to impute zeros in microbiome abundance data using the posterior samples of the underlying true composition. The method is based on an informative prior that has better modeling capability than its predecessors in capturing the essential distributional characteristics of the composition data. Specifically, we propose a BiModal Dirichlet Distribution (BMDD) to model the prior distribution of the true composition. BMDD assumes that each component of the compositional vector follows a bimodal distribution, which is motivated by the observation that some taxa exhibit two modes in their abundance distributions (Lahti et al. 2014). The bimodal distribution could also result from a specific sampling scheme, such as a case-control design, where the cases and controls have different distributions. Moreover, excessive zeros could be efficiently modeled by using a mode sufficiently close to 0 in BMDD. Figure 1a shows the fit of BMDD to the abundance data of four example taxa from the American Gut Project (McDonald et al. 2018). We can see that BMDD provides a better fit than the unimodal Dirichlet distribution. Note that BMDD can also model unimodal distributions and provides a more robust fit to unimodal taxa (see, for example, the top two taxa in Figure 1a) due to its increased modeling capability using more parameters.

**Figure 1:**
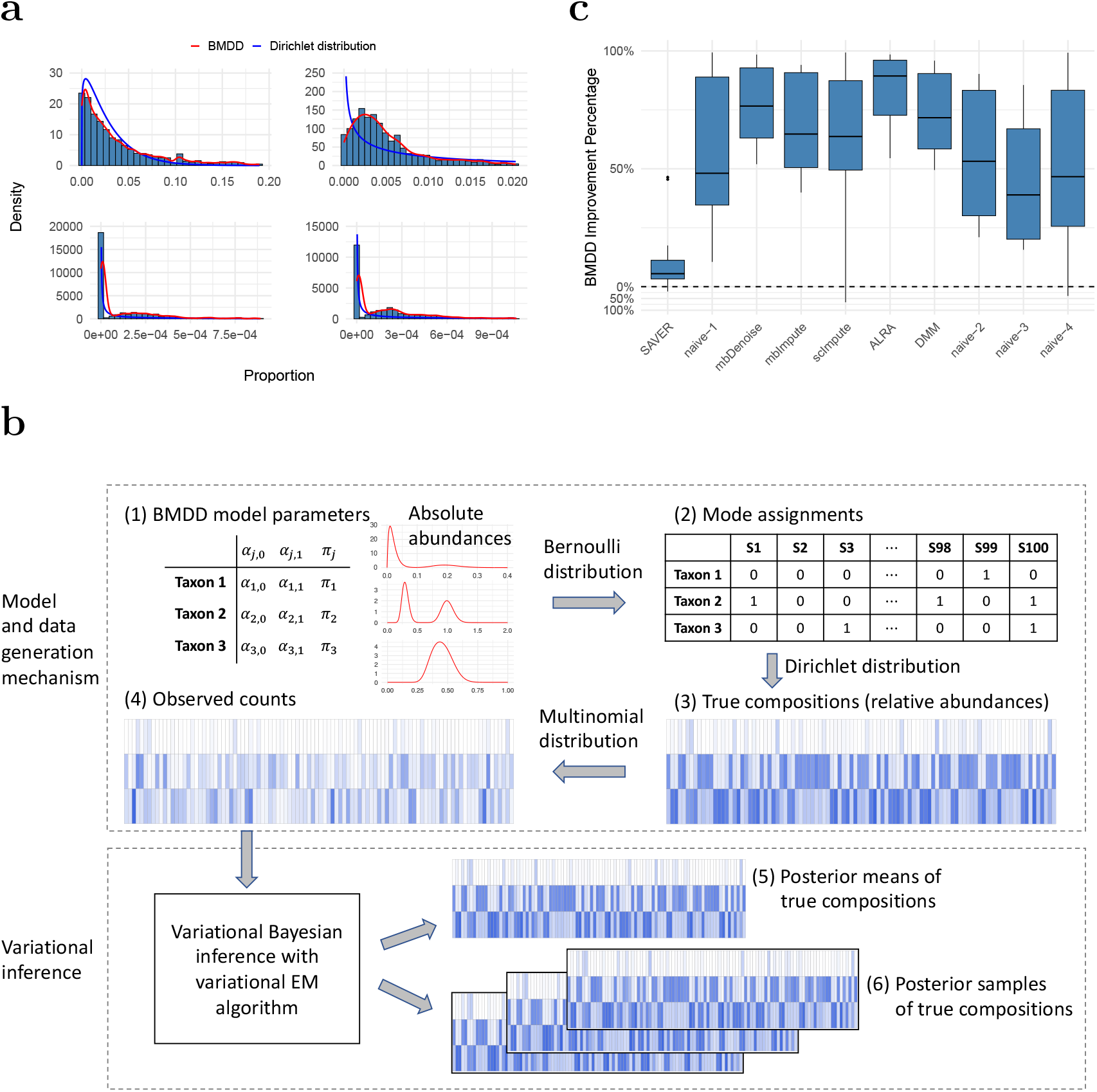
Model fit and imputation performance of BMDD. **a**: BMDD fit on a real human gut microbiome dataset from the American Gut Project. The blue curve is the density of the marginal beta distribution from the fit of the Dirichlet distribution, and the red curve depicts the kernel density estimate based on the posterior means obtained from the fit of BMDD. **b**: Overview of the BMDD. The upper part shows the data generation mechanism of BMDD, and the lower part shows the model computation and inference. Deeper colors indicate higher abundances, and blanks indicate zeros. Details can be found in Section 2.1. **c**: Imputation performance of BMDD compared with competing methods across 15 distance metrics between the estimated and true composition matrices under the simulation setting S1. The performance improvement is expressed as the percentage reduction in distance metrics relative to the respective method.

Our method involves two major steps to find the posterior distribution: (i) we use a mean-field approach to approximate the form of the posterior, which is computationally intractable; (ii) we develop a variational EM algorithm to estimate the hyperparameters in BMDD. The posterior means obtained can be used as estimates of the true compositions. We conducted extensive simulation studies to evaluate the performance of different methods in estimating the true compositions. The results show that BMDD outperforms not only under the correctly specified model but also under misspecified and nonparametric models and thus could be beneficial for downstream statistical tasks such as differential abundance analysis, clustering, and prediction. Based on our model, we propose a multiple imputation approach for differential abundance analysis and demonstrate that it enhances the robustness of log-linear models by accounting for more uncertainty.

## 2 Results

### 2.1 Overview of the BMDD method

Our method is motivated by the observation that some taxa exhibit bimodal abundance distributions (for example, see the bottom two taxa in Figure 1a). One popular model for compositional data, which has a sum-to-one constraint, is the Dirichlet distribution (DD) (Dai et al. 2019). However, DD is not capable of modeling the bimodal distribution observed for some taxa. To address DD’s limitation, we propose the Bimodal Dirichlet Distribution (BMDD) to model the microbiome composition data more flexibly. The BMDD will then be used as the prior distribution under the empirical Bayes framework for zero imputation. To understand BMDD, we first state a basic fact regarding the Dirichlet distribution.

Let *Y*_*j*_ be generated independently of the Gamma distribution with the shape parameter *α*_*j*_ and the scale parameter *ρ*, that is, Gamma(*α*_*j*_, *ρ*) for 1 ≤ *j* ≤ *m*. Setting 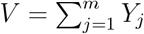, we have (*X*_1_, …, *X*_*m*_) := (*Y*_1_*/V*, …, *Y*_*m*_*/V*) ∼ Dirichlet(*α*_1_, …, *α*_*m*_). *Y*_*j*_, *X*_*j*_ and *V* can be interpreted as the absolute abundance for taxon *j*, relative abundance for taxon *j* and the total microbial load, respectively. Motivated by this basic fact and the intention to capture the bimodal shape of taxa distribution, we consider a two-component mixture distribution for each *Y*_*j*_, that is,

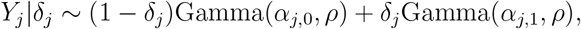

where *δ*_*j*_ ∼ Bernoulli(*π*_*j*_) independently over *j*. Conditional on (*δ*_1_, …, *δ*_*m*_), one has

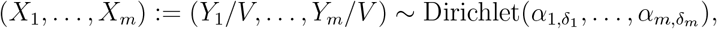

which is independent of the choice of *ρ*. Figure 1b illustrates the BMDD data generation process. Each taxon is parameterized by (*α*_*j*,0_, *α*_*j*,1_, *π*_*j*_), reflecting the mean absolute abundance for the two modes and the probability of the second mode. In Figure 1b(1), we show three representative taxa: one with near zero *α*_*j*,0_ (bimodal with the spike near 0), one with *α*_*j*,0_, *α*_*j*,1_ far apart (bimodal) and one with *α*_*j*,0_, *α*_*j*,1_ in proximity (unimodal). Next, based on the parameter *π*_*j*_, we determine the mode from which the taxon comes for each sample (Figure 1b(2), ‘0’ and ‘1’ represent the two modes). With the mode assignment for each taxon, we generate the true composition (relative abundances) using the Dirichlet distribution (Figure 1b(3)). Note that taxon 1 has many relative abundances near 0. Given the composition and the sequencing depth, we generate the observed counts using a multinomial distribution (Figure 1b(4)). Now, taxon 1 has excessive zeros due to extremely low abundances in most samples.

Under this model, the posterior density of the true composition is computationally intractable, so we use a variational inference approach to approximate the posterior. The hyperparameter of the model is estimated using variational EM. The true compositions are then estimated using the (approximate) posterior mean (Figure 1b(5)). The estimated posterior mean can be used as the imputed composition for downstream statistical analysis and machine learning tasks. Furthermore, we can generate multiple posterior samples from the posterior distribution (Figure 1b(6)). The multiple imputed compositions account for the estimation uncertainty, and the results on individual imputed datasets can be combined to improve the robustness of the analysis using Ruben’s rule (Rubin 2004) or more specialized methods (see Section 2.3 for the example).

### 2.2 True composition estimation for microbiome composition data

The posterior means from our model can be used to estimate the true compositions, upon which downstream statistical analyses can be performed. In this section, we present simulation studies that compare the performance of competing methods to recover the true compositions of microbiome data. We compared our method with mbDenoise and mbImpute, the methods for microbiome data imputation, and also SAVER, scImpute, and ALRA, the methods for scRNA-seq data. Furthermore, we compared our method to a related approach, DMM (Holmes et al. 2012), which uses a Dirichlet multinomial mixture distribution for probabilistic modeling of microbiome data. Unlike BMDD, DMM assumes multiple mixtures of Dirichlet distributions on the full compositional vector rather than on each compositional component. We set the number of Dirichlet mixtures to 10 in our studies. We also included four naive strategies: do nothing (naive–1), add a pseudocount of 1 to all counts (naive–2), replace zeros with 0.5 (naive–3), and impute zeros by 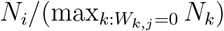 for the *j*th taxon in the *i*th sample (naive–4) as used in Zhou et al. (2022).

To perform a comprehensive evaluation, we considered multiple scenarios. We started by generating data using the BMDD model (setting S1), where the parameters were estimated by fitting BMDD based on the COMBO dataset from a cross-sectional study of the diet effect on stool microbiome composition (Wu et al. 2011). In our evaluation, we simulated *n* = 80 samples and *m* = 100 taxa, reflecting typical microbiome abundance data at the genus level. Based on the true compositions and total counts, multinomial models were used to generate the observed counts.

Denote 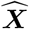 as the estimate of the true composition matrix and 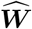 as the imputed count matrix. For our method, we use the posterior mean of the true composition as 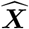. To thoroughly assess the accuracy of different methods in estimating the true composition, we employ fifteen evaluation metrics, which can be broadly divided into three categories: (i) general similarity measures between two composition matrices, including mean squared error, sample-wise distance and taxon-wise distance; (ii) preservation of sample-wise properties, including Shannon’s index, Simpson’s index, Bray-Curtis dissimilarity, Kullback-Leibler divergence, Jensen-Shannon divergence, and Hellinger distance; (iii) preservation of taxon-wise properties, including the Gini coefficient, pair of mean and standard deviation, coefficient of variation, Kolmogorov– Smirnov distance, Wasserstein distance, and pairwise taxon-to-taxon correlation. Smaller values of these metrics indicate better performance. Specific formulations of these metrics are provided in Section A1.1 of the Supplementary Materials.

Figure 1c presents the results of setting S1 in terms of percentage improvement compared to the respective method. Detailed values are provided in Table A1 of the Supplementary Materials. BMDD outperforms competing methods in most evaluation metrics: of the 15 metrics, BMDD ranks first in 12 metrics, second in 2 metrics, and third in the remaining metrics (Table A1). The SAVER method, which considers the count sampling variability and the correlations among taxa, performs very well and achieves the second-best in most of the evaluation metrics. Other methods are substantially less optimal, and oftentimes, the performance could be inferior to the naive methods.

We are also interested in studying the robustness of our method under model misspecification, that is, when the data are not generated according to our BMDD model. As BMDD does not explicitly model the correlations among taxa, we are particularly interested in its performance when the data present such correlations. To achieve this, we conducted experiments in which the data was generated by different models (gamma, log-normal, Poisson, and negative binomial) with correlation structures. We use S2–S5 to denote these settings with the four distributions, respectively. Additionally, we performed simulations where data were generated from non-parametric models, using four real datasets (CDI, IBD, RA, SMOKE) as the templates. We use S6–S9 to denote these non-parametric settings with the four datasets, respectively. Details of these settings can be found in Section A1.2 of the Supplementary Materials.

Under parametric models with correlations (settings S2–S5), BMDD and SAVER continue to outperform other methods (see Figure 2). In general, BMDD outperforms SAVER under the gamma, Poisson, and negative binomial distributions. Under the log-normal distribution, SAVER performs slightly better than BMDD. These results suggest that BMDD is robust to the correlation structure among taxa and model misspecification. Under the nonparametric models (settings S6–S9), the naive methods perform much better than under the parametric settings (Figure 2), which is not unexpected as the observed composition was not much different from the true composition due to the data-generation mechanism. Nevertheless, both the BMDD and the SAVER perform similarly to the naive methods, with BMDD exhibiting some advantages over the SAVER and naive methods. This further highlights the robustness of our BMDD approach.

**Figure 2:**
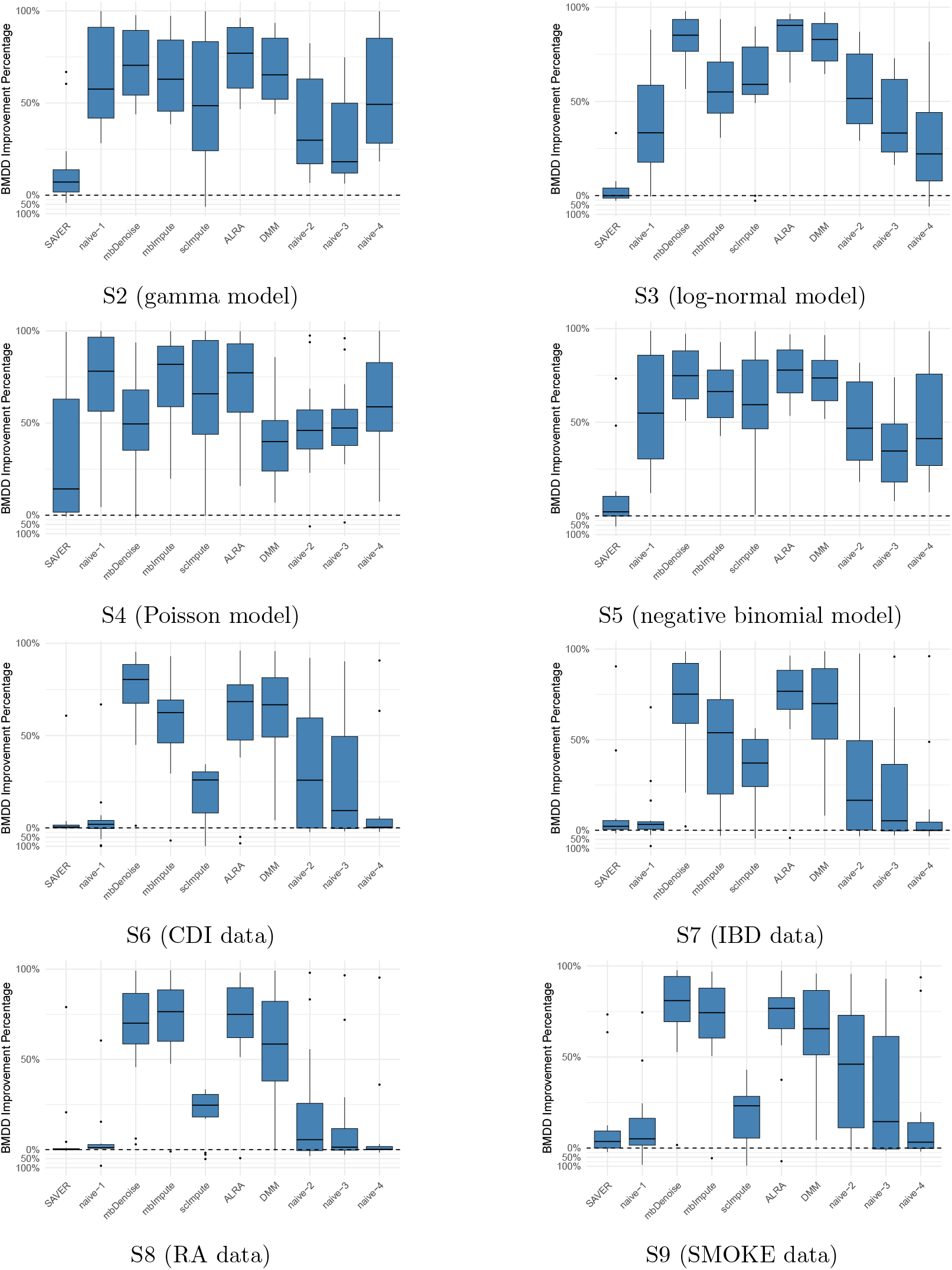
Imputation performance comparison of BMDD with competing methods across 15 distance metrics between the estimated and true composition matrices under settings S2– S9. Formula for the values above zero (the dashed horizontal line): 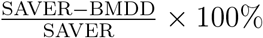, the distance reduction of BMDD relative to SAVER; Formula for the values below zero 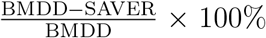, the distance reduction of SAVER relative to BMDD. Likewise for naive-1, mbDenoise, etc.

The BMDD method employs the mean-field variational EM algorithm to estimate the parameters, which is computationally efficient and can be scalable to larger numbers of taxa. Figure 3 shows the execution time (measured in seconds per iteration) of the BMDD algorithm performed on the data generated from the setting S1. We observe that the BMDD algorithm can complete computations (several tens of iterations for most situations) within minutes for moderate sample size (e.g., 200) and number of taxa (e.g., 500) and within hours for a large number of taxa (e.g., 3000).

**Figure 3:**
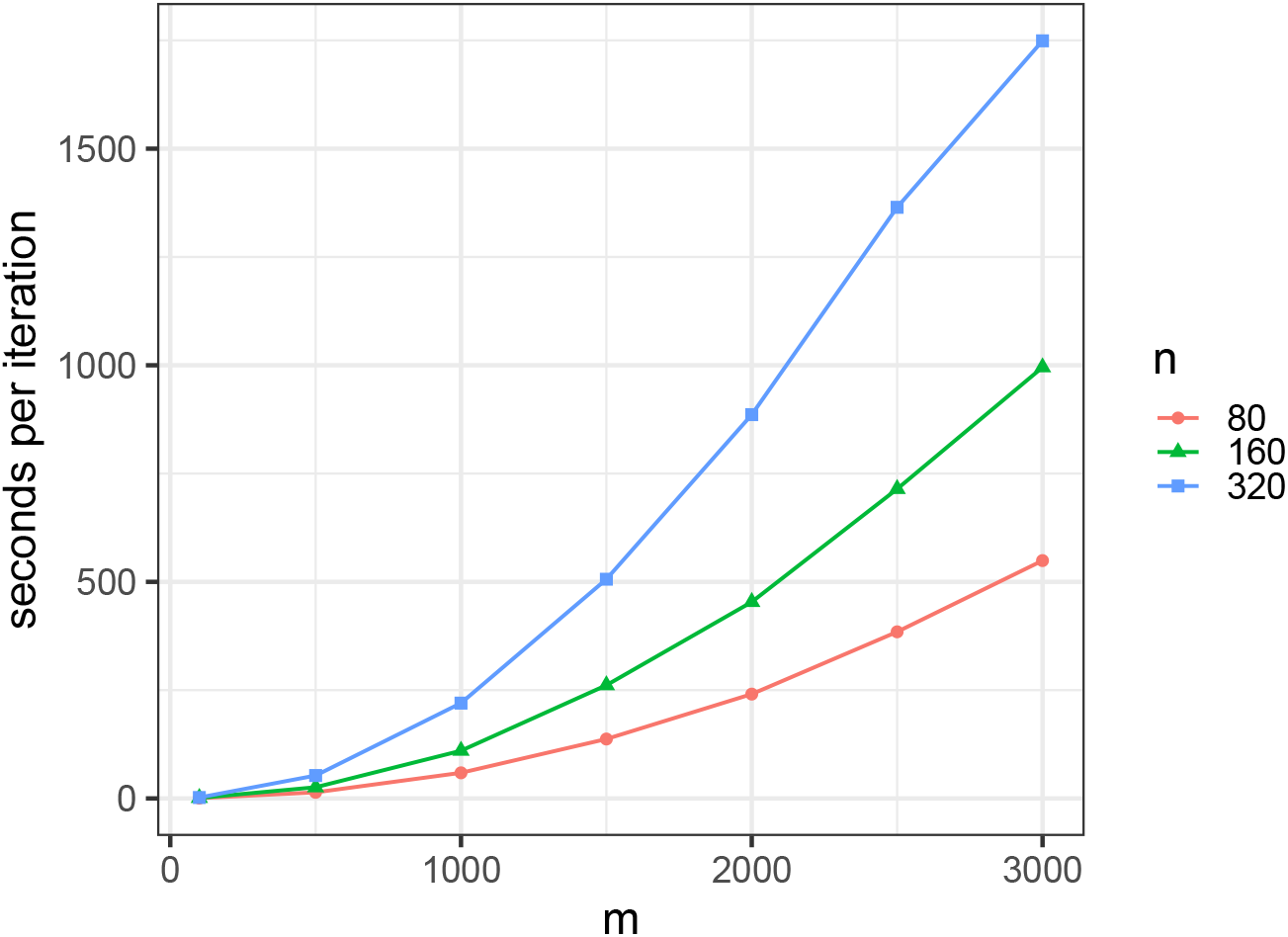
Execution time of the BMDD algorithm (R version 4.3.1 (2023-06-16); Platform: aarch64-apple-darwin20; CPU: Apple M2 Max; Memory: 32 GB). The y-axis represents the time (in seconds) that the BMDD algorithm needs to finish one iteration.

### 2.3 BMDD-based multiple imputation for differential abundance analysis of microbiome data

Differential abundance analysis is a central statistical task of microbiome data analysis, with the aim of identifying microbial taxa whose abundance covaries with a phenotype of interest in medical studies (Yang & Chen 2023). The identified taxa can provide insights into the etiology of the disease and can potentially be used as biomarkers for disease prevention, diagnosis, and treatment (Kashyap et al. 2017). Log-linear models have been widely used in differential abundance analysis of microbiome data due to their simplicity, computational efficiency, and biological interpretability. The recently developed differential abundance analysis tools, ANCOMBC (Lin & Peddada 2020), MaAsLin (Mallick et al. 2021), and LinDA (Zhou et al. 2022), are all based on log-linear models. However, log-linear models require estimating the underlying true compositions and imputing the zeros first. One common approach involves adding a pseudocount, such as 0.5 or 1, to all counts before calculating the composition. The approach is ad hoc, and different pseudocounts could sometimes lead to very different results (Lin & Peddada 2024). As there is much uncertainty in estimating the underlying composition, particularly for those zero counts, a single imputation or a single point estimate may not be sufficient, reducing the robustness of log-linear models. We thus hypothesize that multiple imputations based on posterior samples can significantly enhance the robustness of log-linear models by effectively accounting for uncertainty. Previously, we developed the LinDA method for differential abundance analysis with compositional bias correction (Zhou et al. 2022). One advantage of LinDA is that it can be applied to analyze correlated microbiome data using linear mixed-effects models. As the multiple imputed compositions can be regarded as repeated measurements, we can apply LinDA-LMM conveniently to integrate multiple imputations, thus accounting for the uncertainty in composition estimation. In the following, we conduct simulation studies and real data applications to compare the performance of original LinDA (which uses the pseudocount approach), LinDA with the posterior samples from our method (denoted as LinDA-BMDD), and LinDA with the posterior samples from SAVER (denoted as LinDA-SAVER) in terms of FDR control and power.

We employed similar simulation setups as those used in the LinDA paper(Zhou et al. 2022). Specifically, we included the setups with a log-normal distribution for absolute abundance, a gamma distribution for absolute abundance, and a negative binomial distribution for observed abundance (setups S0, S3, and S7 in Zhou et al. (2022), respectively). In addition, we added a setup with Poisson distribution for the observed abundance in this study. Details of data generation can be found in Section A2.1 in the Supplementary Materials. We considered different configurations for the number of taxa and sample size: *m* = 50, 200, 500 and *n* = 50, 200. We adopted a moderate signal density (10% differential taxa) and signal strength such that we could see the power difference between methods.

We used 100 posterior samples of the true composition matrix for both LinDA-BMDD and LinDA-SAVER, so the input sample size for these two methods was 100*n*. Under the gamma model, LinDA-BMDD controls the FDR across settings, with nonsignificant FDR inflation when the number of taxa is small (Figure 4a). In contrast, LinDA and LinDA-SAVER suffer from more serious FDR inflation in many settings, especially when the number of taxa is small. In terms of power, LinDA-BMDD has a slight power loss compared to the original LinDA, but the loss is not substantial, indicating that the increased robustness retains most of the statistical power. Under the log-normal model (Figure 5), LinDA-SAVER has the worst FDR control, with inflation worsening as the number of taxa increases. LinDA and LinDA-BMDD control the FDR overall, with only slight FDR inflation when the number of taxa is small. Under the Poisson and negative binomial (Figure 5), LinDA-BMDD again shows much better FDR control than LinDA-SAVER and LinDA.

**Figure 4:**
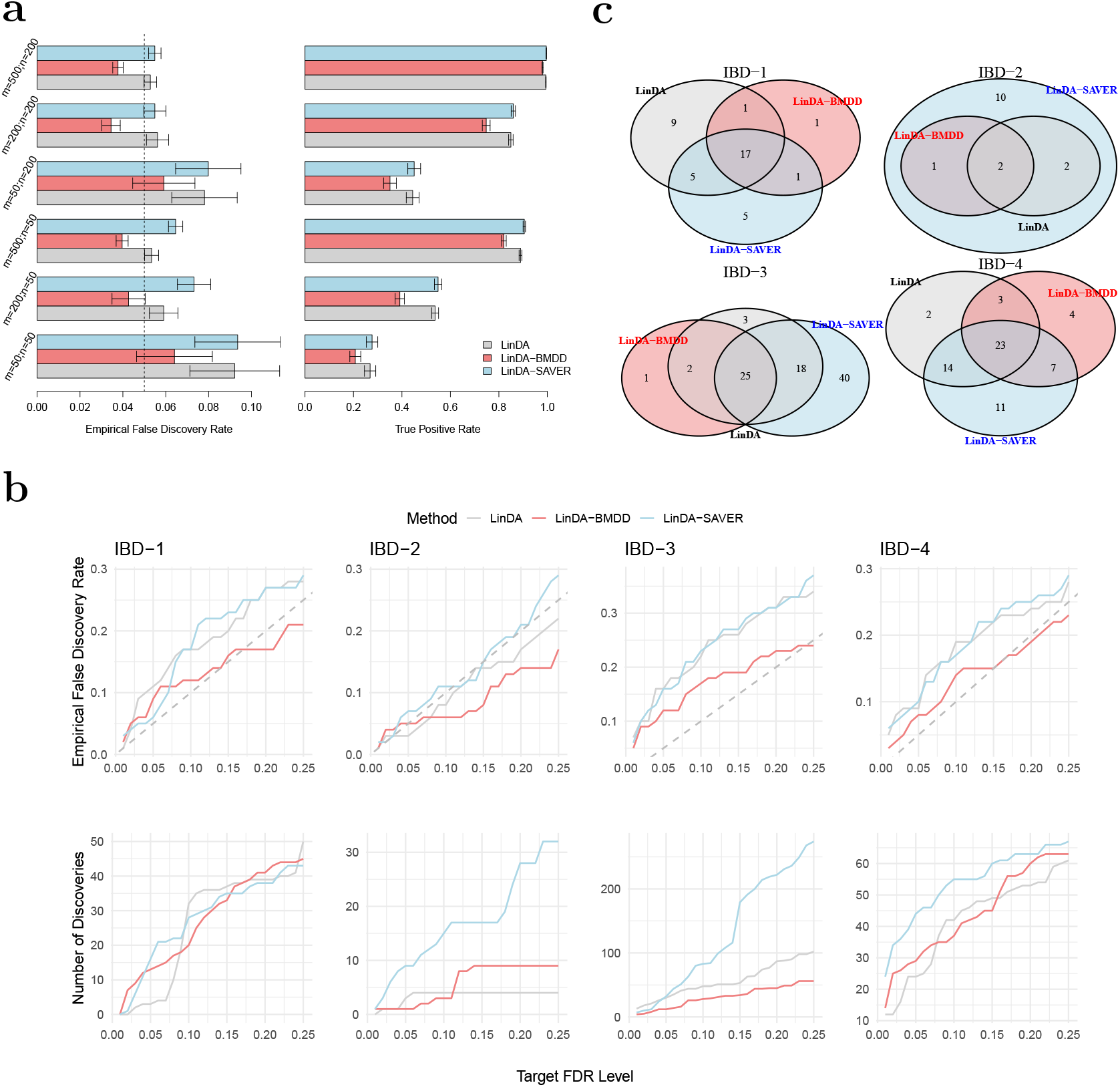
Performance of BMDD in differential abundance analysis of microbiome data. **a**: Simulation results for differential abundance analysis under the gamma model. Empirical false discovery rates and true positive rates were averaged over 500 simulation runs. Error bars represent the 95% confidence intervals and the dashed vertical line indicates the target FDR level of 0.05. **b**: (Bottom) Number of discoveries vs. target FDR level for the real datasets; (Top) Empirical FDR vs. target FDR level for the shuffled real datasets. The dashed gray line represents the diagonal. The results were averaged over 100 simulation runs. **c**: Overlaps of differential taxa with target FDR level of 0.1 for the four real datasets, IBD-1, IBD-2, IBD-3 and IBD-4.

**Figure 5:**
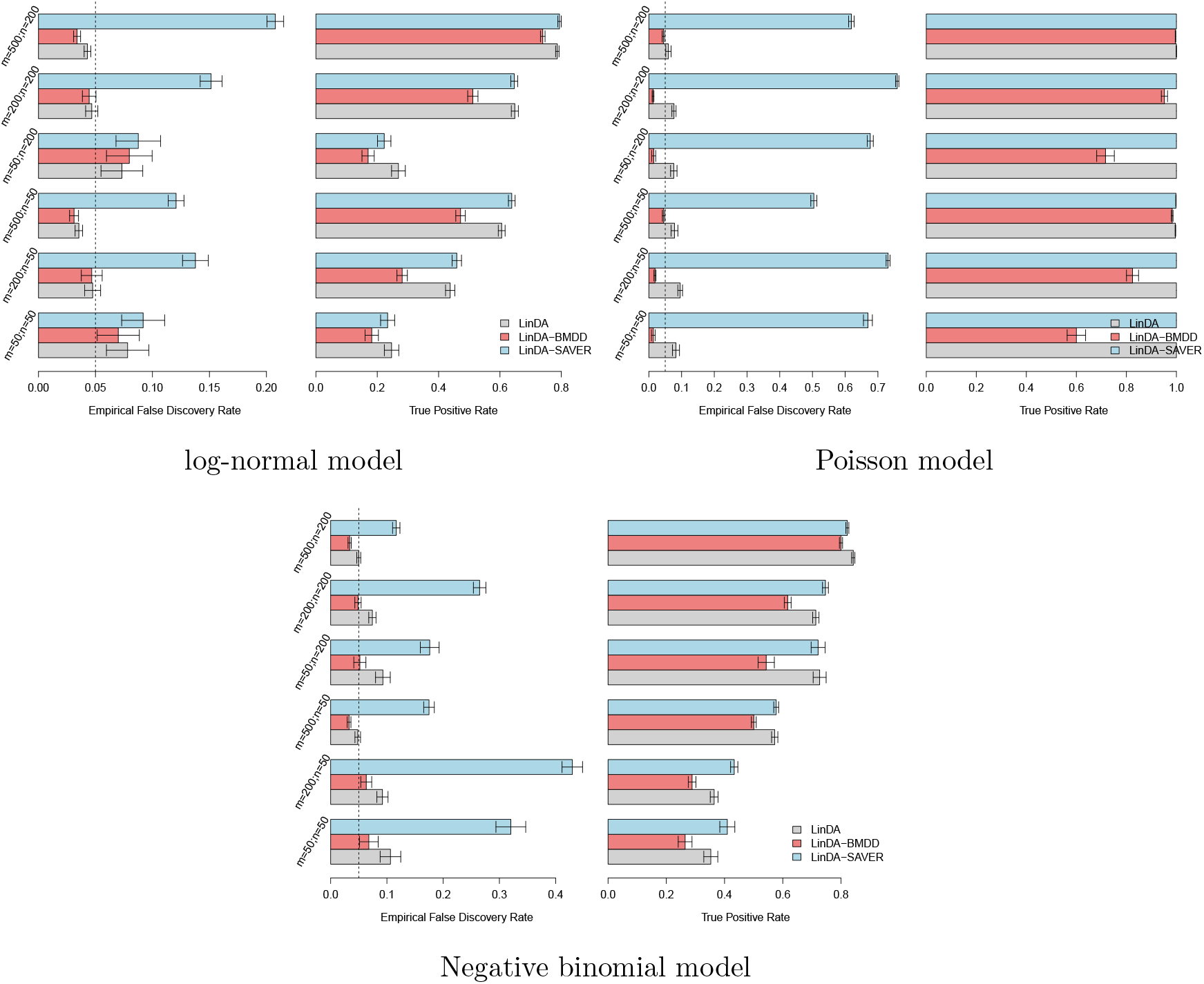
Simulation results for differential abundance analysis under the log-normal, Poisson, and negative-binomial models, respectively. Error bars represent the 95% confidence intervals and the dashed vertical line indicates the target FDR level of 0.05.

We reason that the inferior FDR control by LinDA-SAVER could be due to “information diffusion,” as zeros were imputed based on information from other taxa based on the underlying correlation structure. When there are many differential taxa, the differential signals may diffuse to those non-differential ones, increasing the false positive rate. Therefore, applying LinDA-SAVER is not recommended for differential abundance analysis, even if it is apparently highly powerful in some settings.

Overall, when the model is correctly specified for LinDA (that is, log-normal), the original LinDA performs well, and there is no significant benefit in FDR control with the use of the multiple imputation approach. However, when the model is misspecified, LinDA with BMDD imputation significantly reduces the FDR inflation with only slight power loss. These simulation studies confirm that integrating the posterior inference using BMDD in LinDA leads to a more robust testing procedure by taking into account the uncertainty of the underlying true abundance under zero counts.

Next, we applied the three competing methods to four real datasets from case-control gut microbiome studies of inflammatory bowel disease (IBD). The four genus-level abundance datasets were retrieved from the MicrobiomeHD database (Duvallet et al. 2017). We excluded samples with fewer than 1000 read counts and taxa, which appear in less than 20% of the samples. The characteristics of the filtered datasets are shown in Table A2 in Section A2.2 of the Supplementary Materials. We compared the number of discoveries of LinDA, LinDA-BMDD, and LinDA-SAVER at different FDR levels (0.01–0.25) and studied their overlap patterns at the target FDR of 0.1. As the ground truth is unknown, the assessment of the FDR control performance is difficult. However, we could at least evaluate the FDR control under the global null, where we disrupt the signals by shuffling the group labels. The empirical FDR was then calculated as the percentage of repetitions (random group label shuffling) that made any discoveries. In all analyses, we used winsorization at quantile 0.97 to reduce the impact of potential outliers as recommended in Chen et al. (2018).

From Figure 4b, we can see that, in general, the patterns are consistent with the simulation studies. LinDA-BMDD significantly improves the FDR control over the original LinDA and LinDA-SAVER. The power of LinDA-BMDD is comparable to or slightly less than the original LinDA, echoing the simulation results. LinDA-SAVER has substantially more power than LinDA-BMDD for two datasets (IBD-2 and IBD-3), but the increased power could be due to its inefficient FDR control due to “information diffusion” discussed in the simulation results.

The overlap analysis (Figure 4c) reveals that most of the discoveries made by LinDA-BMDD at 10% FDR were also recovered by LinDA and/or LinDA-SAVER. In contrast, many discoveries made by LinDA-SAVER were not found by the other two methods. Those taxa recovered by LinDA-BMDD were also supported by previous literature. For example, in IBD-1, all three methods made similar numbers of discoveries, while only LinDA-BMDD offered adequate FDR control under the global null based on shuffled datasets. At 10% FDR, LinDA-BMDD, LinDA-SAVER, and LinDA identified 20, 28, and 32 differential taxa, respectively. The differential taxa identified by LinDA-BMDD showed a large overlap with LinDA-SAVER and LinDA. Nineteen taxa were also identified by either or both of the other methods. Only one taxon was unique to LinDA-BMDD. In contrast, LinDA-SAVER and LinDA had 9 and 5 method-specific taxa. The taxon identified only by LinDA-BMDD belonged to Clostridium XlVa of the Lachnospiraceae family. Lachnospiraceae, including Clostridium clusters IV and XIVa, were previously implicated in IBD (Atarashi et al. 2013). Detailed studies of the results for IBD-2, IBD-3 and IBD-4 can be found in Section A2.3 in the Supplementary Materials.

To see if the increased robustness of differential abundance analysis is limited to LinDA, we further tested BMDD on ANCOM-BC (Lin & Peddada 2020, 2024), one of the most popular differential abundance analysis methods. We compared the performance of the original ANCOM-BC to ANCOMBC-BMDD and ANCOMBC-SAVAER. Since ANCOMBC is computationally much slower, we used 20 posterior samples of the true composition matrix for ANCOMBC-BMDD and ANCOMBC-SAVER. The results on the number of discoveries and empirical FDR at different FDR levels are presented in Figure A1 in the Supplementary Materials. We observed that the original ANCOM-BC failed to control the FDR effectively: its empirical FDRs across different datasets and FDR levels were almost equal to 1, indicating that it always made some rejections when the group labels were shuffled. Interestingly, with the multiple imputation approach, using either the posterior samples from BMDD or SAVER, the FDRs were conservatively controlled. We note that in the newer version ANCOM-BC (ANCOM-BC2) (Lin & Peddada 2024), the authors recommended a sensitivity analysis based on different pseudocount additions. Using this strategy, the authors demonstrated significantly improved robustness of ANCOM-BC with more accurate FDR control. In comparison, our method proposed a more principled way to handle zeros without the need for sensitivity analysis.

## 3 Discussion

In this study, we proposed a Bayesian approach to estimate the true composition of the microbiome from the observed sequencing count data. We modeled the underlying true abundance using a mixture of gamma distributions for each taxonomic component, leading to a bimodal Dirichlet distribution for the taxonomic compositions. Using mean-field approximations for posteriors, we developed an efficient variational EM algorithm to obtain the estimates. Our results show that BMDD provides the best overall estimate of true compositions compared to existing methods for microbiome data based on extensive simulations. We also show that the proposed multiple imputation approach using posterior samples significantly improves the robustness of differential abundance analysis methods such as LinDA and ANCOM-BC.

The SAVER method, developed for single-cell sequencing data, also shows competitive performance. However, the constructions of the two methods are very different. SAVER employs a Poisson distribution with a gamma prior to modeling the counts and explores the relationship between genes in the same cell/sample (pooling information across genes), while BMDD utilizes a gamma mixture model for the counts of each taxon (pooling information across samples) and operates on the resulting bimodal Dirichlet distribution for the compositions. Thus, it is possible to combine the strengths of both methods to further improve the imputation performance. However, in differential abundance analysis, borrowing information from correlated taxa can lead to inflated false positive rates due to “information diffusion” from those truly differential taxa.

One drawback of BMDD is its limited ability to model the inter-taxa correlations. However, since the BMDD-based imputation method mainly borrows information from the abundance distribution of the same taxon in other samples, ignoring correlations among taxa is not expected to seriously affect its imputation performance. As suggested by the simulation results in Figure 2 and Figure 5, BMDD model appears robust to the correlation structure among taxa.

In addition to differential abundance analysis methods, we envision that multiple imputation using BMDD will be useful in other application settings such as community-wide testing, clustering, and prediction (Zhang et al. 2022, Shi et al. 2022, 2016). How to integrate multiple imputations efficiently in these statistical tasks warrants further investigation. Although our method was motivated by the analysis of microbiome data, it can be applied in principle to other zero-inflated compositional data analyses, such as scRNA-seq data. However, rigorous benchmarking studies must be performed before BMDD can be recommended for other applications.

Finally, our method can be potentially extended to accommodate both sample covariates and phylogeny among taxa, as outlined in Section A3 of the Supplementary Materials, to further improve the efficiency of the imputation. As this adds another layer of complexity, more computationally efficient methods are needed. Moreover, its robustness in the presence of misspecified covariates or incorrect phylogeny requires further study. We leave this as our future research direction.

## 4 Materials and Methods

We develop a new probabilistic model for zero-inflated microbiome data. In particular, we assume the observed counts are generated from a multinomial distribution with the true composition following a BMDD to capture the variation of the composition data.

### 4.1 Definition of BMDD

We first state a basic fact regarding the Dirichlet distribution, which inspires our definition of BMDD. Let *Y*_*j*_ be generated independently from the Gamma distribution with the shape parameter *α*_*j*_ and scale parameter *ρ*, that is, Gamma(*α*_*j*_, *ρ*) for 1 ≤ *j* ≤ *m*. Setting 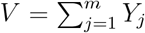, we have (*Y*_1_*/V*, …, *Y*_*m*_*/V*) ∼ Dirichlet(*α*_1_, …, *α*_*m*_). Motivated by this basic fact and the intention to capture the bimodal shape of taxa distribution, we consider a two-component mixture distribution for each *Y*_*j*_, that is,

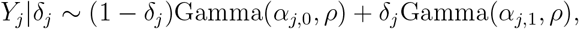

where *δ*_*j*_ ∼ Bernoulli(*π*_*j*_) independently over *j*. Conditional on (*δ*_1_, …, *δ*_*m*_), one has

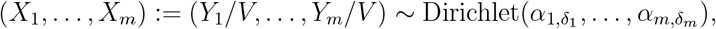

with *X*_*j*_ = *Y*_*j*_*/V*, which is independent of the choice of *ρ*. We name the above probability model the BiModal Dirichlet Distribution (BMDD). An alternative way to define BMDD is through the following hierarchical model:

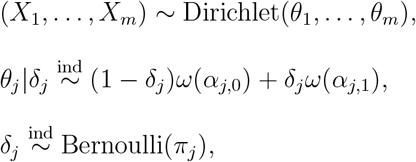

where *ω*(*x*) denotes a point mass at *x*.

### 4.2 The probabilistic model

Let *W*_*i,j*_ represent the observed read count of taxon *j* in individual *i*. For the *i*th individual, the total count of all taxa, *N*_*i*_, is determined by the sequencing depth and DNA materials. Let ***X***_*i*_ = (*X*_*i*,1_, …, *X*_*i,m*_) denote the unobserved true composition for the *i*th individual. Given *N*_*i*_, it is natural to model the stratified count data over *m* taxa as a multinomial distribution

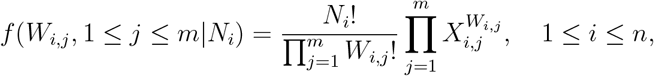

where 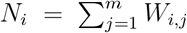. Assuming that 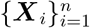 are i.i.d samples generated from the BMDD, then we have the following hierarchical model:

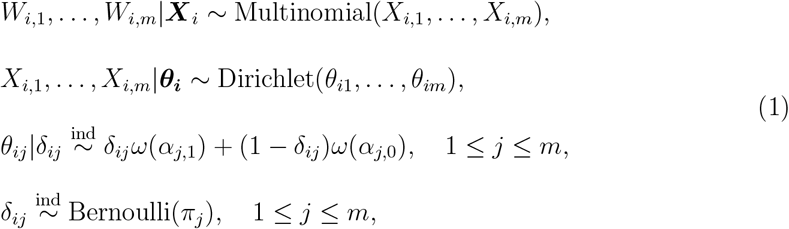

where ***θ***_***i***_ = (*θ*_*i*1_, …, *θ*_*im*_). The goal here is to estimate the hyperparameters (***π, α***) in the BMDD as well as the posterior distribution of ***X*** with ***X*** = (***X***_1_, …, ***X***_*n*_), ***π*** = (*π*_1_, …, *π*_*m*_) and ***α*** = (*α*_*j,k*_)_1*≤j≤m,k*=0,1_.

#### Remark 1

Our model (1) can be extended to incorporate optional metadata, including sample covariates and taxon phylogeny. The details can be found in Section A5 of the Supplementary Materials.

### 4.3 Variational inference

Let ***W*** = (***W*** _1_, …, ***W*** _*n*_) with ***W*** _*i*_ = (*W*_*i*,1_, …, *W*_*i,m*_). The joint likelihood of (***W***, ***X***) is given by

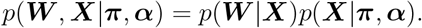

The major challenge is the computational intractability of the density

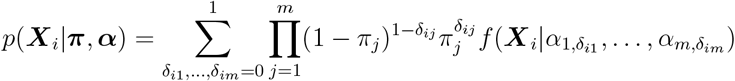

with *f* (***x***|*α*_1_, …, *α*_*m*_) denoting the density of Dirichlet(*α*_1_, …, *α*_*m*_).

To overcome this difficulty, we consider the variation frequentist estimate based on the mean-field approximation for the posterior. Specifically, we consider a class of distributions of the form

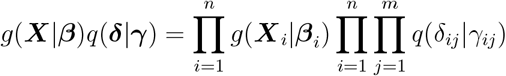

where ***δ*** = (*δ*_*ij*_)_1*≤i≤n*,1*≤j≤m*_,***γ*** = (*γ*_*ij*_)_1*≤i≤n*,1*≤j≤m*_ with 0 ≤ *γ*_*ij*_ ≤ 1, ***β*** = (***β***_1_, …, ***β***_*n*_) with ***β***_*i*_ = (*β*_*i*1_, …, *β*_*im*_), *g*(·|***β***_*i*_) represents the density function of Dirichlet(*β*_*i*1_, …, *β*_*im*_) and *q*(·|*γ*) denotes the probability mass function of Bernoulli(*γ*). Here *g*(***X***|***β***)*q*(***δ***|***γ***) is a variational distribution that serves as a surrogate for the posterior distribution *p*(***X, δ***|***W***, ***π, α***). As seen from the derivations below, the form of the variational distribution leads to a computationally feasible coordinate ascent algorithm.

To motivate our algorithm, we note that

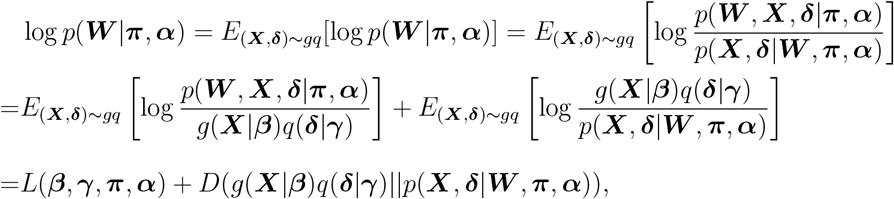

where *L*(***β, γ, π, α***) is called the evidence lower bound (ELBO) in variational Bayes, and *D*(·∥·) is the Kullback-Leibler (KL) divergence of the approximation from the true posterior. Therefore, maximizing the ELBO is equivalent to minimizing the KL divergence. We propose a variational EM algorithm to estimate the unknown parameters, which involves iterations of the following two steps:

1. E-step: fixing (***α, π***), update the parameter (***β, γ***) in the mean field approximation through the coordinate ascent mean-field variational inference (Bishop 2006);
2. M-step: fixing (***β, γ***), update the hyperparameters (***α, π***) by maximizing the ELBO. Below, we describe these two steps in detail.

### 4.4 Variational EM algorithm

#### E-step: update the mean-field approximation

Given ***π*** and ***α***, we aim to update the mean-field distribution by maximizing the ELBO. We employ one of the most commonly used algorithms for solving this optimization problem, the coordinate ascent variational inference (CAVI). According to CAVI, we deduce the optimal form of *g*(***X***_*i*_|***β***_*i*_),

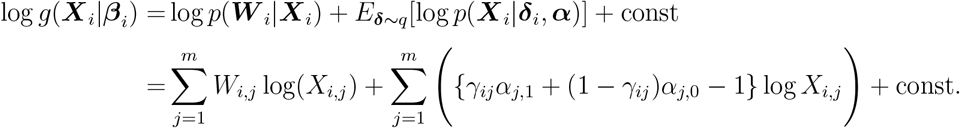

It thus implies that *β*_*ij*_ = *W*_*i,j*_ + *γ*_*ij*_*α*_*j*,1_ + (1 − *γ*_*ij*_)*α*_*j*,0_ for 1 ≤ *j* ≤ *m*. On the other hand, the optimal form of *q*(*δ*_*ij*_ |*γ*_*ij*_) is given by

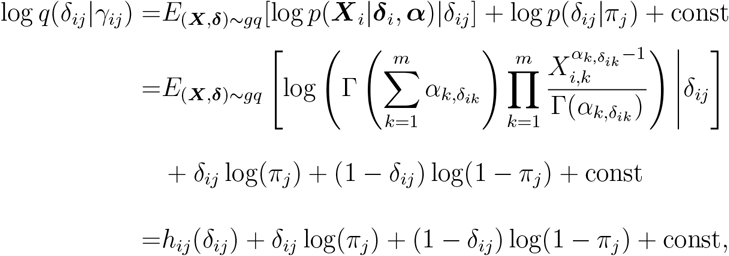

where 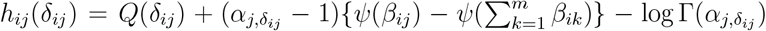 with 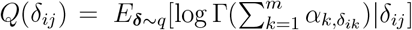, and we have used the fact that 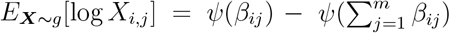, where *ψ* denotes the digamma function. Therefore, we obtain

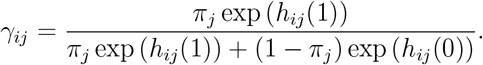

One can approximate the expectation in *Q*(*δ*_*ij*_) via the Monte Carlo approach. Note that *h*_*ij*_ depends on *γ*_*ik*_ for *k*≠ *j*. One needs to iterate the above updates across *j* = 1, 2, …, *m* until convergence. The iteration can be done in a parallel fashion across *i* = 1, 2, …, *n*.

##### Remark 2

The Monte Carlo approximation for *Q*(*δ*_*ij*_) can be computationally very intensive. To reduce the computation cost, we consider the following approximate approach.

Using the fact that the log gamma function is convex on the positive real line, we have

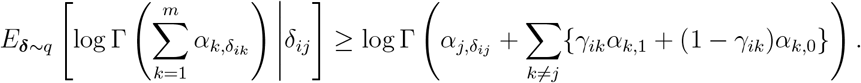

Maximizing this lower bound leads to the optimal form of *q* given by *h*_*ij*_ with the term *Q*(*δ*_*ij*_) in *h*_*ij*_ being replaced by 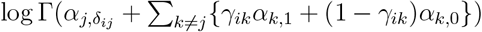. This approach is computationally much faster than the Monte Carlo approximation.

#### M-step: update the hyperparameters

Next, we update the hyperparameters by optimizing the ELBO with respect to (***π, α***):

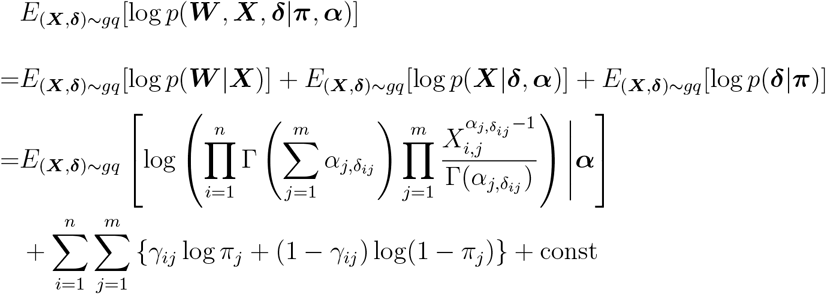

Again the term 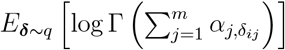 is not tractable. We address this issue using the same trick as in Remark 2 and propose to update ***π*** by letting 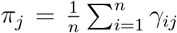 and update ***α*** by maximizing the following objective function through the optimization function “nlminb” in the R software,

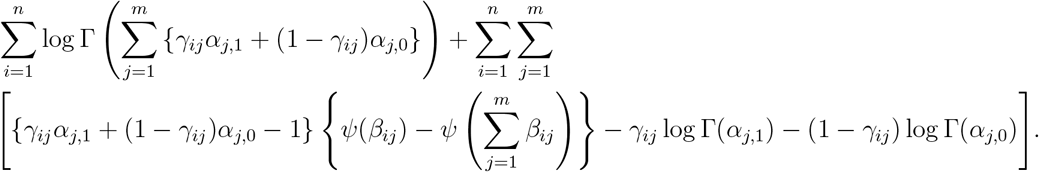

## Declarations

### Ethics approval and consent to participate

Not applicable. The work uses public datasets.

### Consent for publication

Not applicable. The manuscript contains no individual person’s data.

### Funding

The work is supported by National Natural Science Foundation of China 12201384, Fundamental Research Funds of Shanghai University of Finance and Economics, Research Funds of the School of Statistics and Data Science at Shanghai University of Finance and Economics (Zhou), National Institute of Health R01GM144351 (Chen & Zhang), National Science Foundation DMS1830392, DMS2113359, DMS1811747 (Zhang), National Science Foundation DMS2113360, and Mayo Clinic Center for Individualized Medicine (Chen).

### Availability of data and materials

The Supplementary Materials provide the technical details and additional application results referenced in the main text. The BMDD package is available at GitHub (https://github.com/zhouhj1994/BMDD).

### Competing interests

The authors declare that they have no competing interests.

### Authors’ contributions

XZ and JC conceived, designed, and supervised the study. XZ and HZ developed the algorithm and implemented the software. HZ performed the evaluations and led the manuscript writing. All authors contributed to the revision of the manuscript and approved the final manuscript.

## Acknowledgments

Not applicable.

## Notes

### Competing Interest Statement

The authors have declared no competing interest.

